# A consensus view on the folding mechanism of protein G, L and their mutants

**DOI:** 10.1101/2022.04.07.487494

**Authors:** Liwei Chang, Alberto Perez

**Affiliations:** Department of Chemistry, University of Florida, Gainesville, FL 32611, USA; Quantum Theory Project, University of Florida, Gainesville, FL 32611, USA

## Abstract

Much of our understanding of folding mechanisms comes from interpretations of experimental ϕ and ψ value analysis – relating the differences in stability of the transition state ensemble (TSE) and folded state. We introduce a unified approach combining simulations and Bayesian inference to provide atomistic detail for the folding mechanism of protein G, L and their mutants. Protein G and L fold to similar topologies despite low sequence similarity, but differ in their folding pathways. A fast folding redesign of protein G, NuG2, switches folding pathways and folds through a similar pathway with protein L. A redesign of protein L also leads to faster folding, respecting the original folding pathway. Our Bayesian inference approach starts from the same *prior* on all systems and correctly identifies the folding mechanism for each of the four proteins – a success of the force field and sampling strategy. The approach is computationally efficient and correctly identifies the TSE and intermediate structures along the folding pathway in good agreement with experiments. We complement our findings by using two orthogonal approaches that differ in computational cost and interpretability. Adaptive sampling MD combined with Markov State Model provide a kinetic model that confirms the more complex folding mechanism of protein G and its mutant. Finally, a novel fragment decomposition approach using AlphaFold identifies preferences for secondary structure element combinations that follows the order of events observed in the folding pathways.

## Introduction

Machine Learning approaches have recently revolutionized structural biology by predicting accurate 3D structures of proteins at atomistic resolution from their sequence with high confidence and accuracy^1,2^. In an unprecedented effort, massive structural libraries for different organisms have been freely released to the community based on AlphaFold (AF) predictions^3^. These advances are further leading to novel protein designs based on the idealized versions of proteins the model has learned^4^. The protein folding problem refers to cracking two codes: (1) what is the structure of a protein based on the sequence, and (2) what is the process by which proteins fold so fast. While the first code has now been mostly solved through machine learning, the answer to the second code remains unclear. Solving the second code refers to identifying the pathways a protein takes that allow it to fold much faster than a random search among all possible configurations according to Levinthal’s paradox^5^. Experimental data is too sparse for training a neural network and even when data is available, the mapping between experimental and atomistic interpretations of the folding pathways are challenging. Computational approaches using simulations are often used to complement experimental pathways with atomistic insight, but such simulations tend to be computationally demanding and of difficult interpretation.

In this work, we revisit the folding mechanisms of protein G and protein L, two proteins with low sequence similarity (~15%) that fold into the same topology composed of a central-helix packed against two antisymmetric β-hairpins. These two proteins have been the object of experimental^6–12^ and computational^13–22^ studies to determine their folding routes, and have led to the design of faster folding mutants. Mutant NuG2 replaces 11 amino acids that form a native type I turn in the N-terminal hairpin of protein G with a thermodynamically favorable type I’ turn – which also results in faster folding kinetics^23^. For protein L, design efforts focus on the C-terminal hairpin, where the turn contains three consecutive residues with positive backbone ϕ angles that likely caused low intrinsic stability for this hairpin. While the kinetics are improved ~20 fold with respect to protein L, the stability is not increased^13^.

Initial folding kinetics studies suggested a two-state folding pattern for protein G^6^, while experiments using stopped-flow fluorescence methods and thermal unfolding simulations demonstrated the presence of an on-pathway intermediate state^16^. More recently, experiments combining multiple probes on the submillisecond timescale and large-scale simulation analysis^8^ have led to a more complex folding mechanism with multiple intermediates. Initial studies pointed to the formation of the C-terminal hairpin as the rate-limiting step. Redesigning the N-terminal hairpin leads to a higher stability hairpin (NuG2 mutant) that induces a switch in the folding pathway as well as the folding rate^24^.

Despite the slower folding rate of protein L, its folding pathway is simpler than that of protein G, with only the N-terminal hairpin involved in the transition state ensemble (TSE) according to mutational ϕ analysis^9^. More recent ψ analysis experiments showed that the a non-native C-terminal hairpin was also established in the TSE^10^. Despite their small size, characterizing the folding mechanism of protein G and L involves considerable challenges as multiple factors can influence experimental observations without revealing the underlying atomistic detail^7^.

Molecular dynamics (MD) simulations are a promising computational tool to bridge between experiments and atomistic interpretation of folding pathways. In practice, the approach is limited due to the disparity between accessible simulation times (millisecond timescale) and the slow timescale of the folding pathways (microsecond to second folders for the proteins in this study). Millisecond long simulations on a set of 12 structurally diverse, fast-folding proteins remain the golden standard in the field^25^. One of these proteins, a triple mutant of NuG2 was simulated at its melting temperature for more than 1 ms, sampling multiple folding and unfolding events. Even with these ensembles, sampling of the possible unfolded states and pathways remains incomplete.^26^ Thus, sampling methods such as replica exchange molecular dynamics (REMD),^27^ adaptive sampling,^28^ path based sampling^29^ and metadynamics^30^ to name a few, are often used to enhance the sampling efficiency of MD simulations. However, no study has provided a unified, atomistic level comparison and interpretation of the folding behaviors of protein G, L and their mutants. Most computational work has focused on the faster folding variants NuG2 (microsecond folders) as it is computationally more affordable compared to the slower folding time of protein G^31^.

In this work we provide a unified view of the folding pathway preference for the four proteins by accelerating molecular dynamics with Bayesian inference using minimal assumption from ψ and ϕ analysis, extensive adaptive sampling with Markov State Model (MSM) analysis, and the state-of-the-art protein structure prediction method AlphaFold (AF). Feeding the same information to the four different sequences studied here, the method successfully distinguishes folding pathways for each sequence and finds metastable states compatible with the literature (e.g., out of registry β-strand pairings).

## Results

### Accelerating the nucleation of early folding events leads to global folding

Previous studies have indicated the formation of an initial hairpin plays a crucial role in the folding process of protein G and L, but whether both hairpins or only one appear in the folding transition state differs between experimental ϕ and ψ analyses^6,7,13^. We introduce data to bias conformations of the known β-strands on extended conformations (see methods), leading to more efficient nucleation events when two strands come in proximity to each other (Fig.1b). Although we introduce equivalent data for the four strands, we require that only two strands satisfy the data in simulations using Bayesian inference with our Modeling Employing Limited Data (MELD) approach^32,33^ (Fig.1b). By biasing the conformations of individual strands, all possible combinations (and orientations) of strand pairings are possible. We evaluate two key questions from the generated ensembles starting from fully extended conformations of protein G, L and their mutants: (1) whether the designed sampling strategy combined with the physics model is capable of folding proteins to their native structure; (2) whether the identified possible intermediates and folding pathways agree with experiment observation.

**Figure 1.**
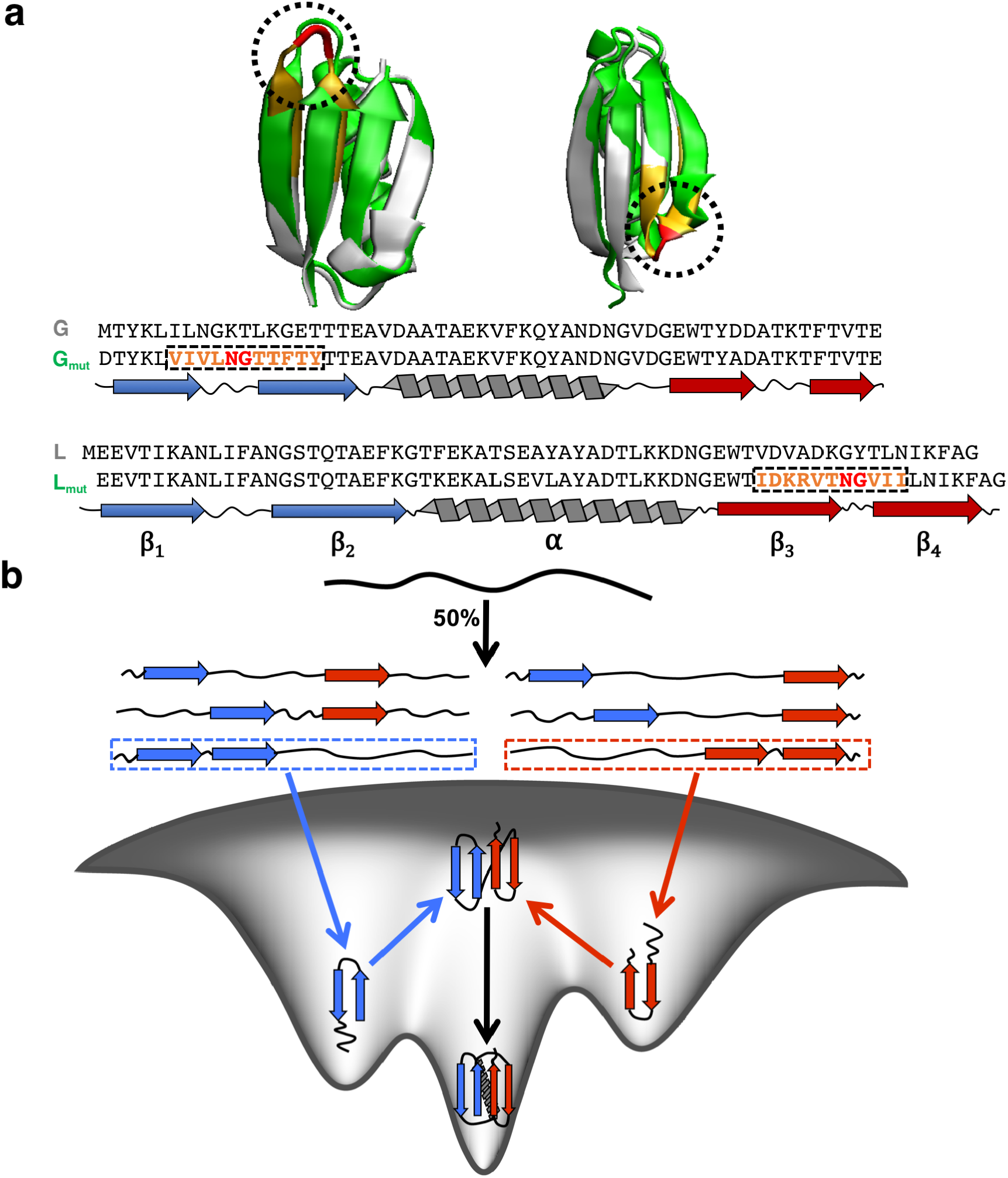
Native structures and designed MELD simulation protocol. **a** Native structures of wild type (white) protein G (PDB: 3GB1), protein L (PDB: 1HZ5), and the mutant (green) of protein G (PDB: 1MI0), protein L (PDB: 1KH0) with their corresponding sequence and secondary structure assignments. Mutation regions are colored in orange and red for both structures and sequences. **b** Illustrating designed protocol for folding simulations of proteins considered here in MELD. Selective secondary structure restraints on four beta strands are enforced, where N/C-terminal hairpin is favored based on conformations generated from Amber force field.

We first find that the designed sampling protocol is able to sample native folds for all proteins and identify the native state based on clustering in excellent agreement with the experimental structure. (Fig. S1) All four proteins reached conformations with Cα RMSD lower than 2 Å and a native contact fraction higher than 0.9. Previous simulations using heuristics guiding to the end state^33^ require short simulations (1*μ*s of sampling, 30 replicas) to systematically fold proteins G and L with high populations. The current simulation protocol requires much longer sampling times, with ~8-20 *μ*s and 30 replicas needed for each system. There is a surprising difference in behavior between protein G/G_mut_ and L/L_mut_: in the latter pair, multiple independent “walkers” are able to identify the native state, whereas in the former pair few “walkers” find the native state (Fig. S2 and S3). This observation is consistent with the higher complexity of protein G folding pathways, where protein G and its mutant struggled to progress to folding transition state compared with protein L and its mutant. We describe below the folding mechanisms identified from our simulations.

### A competitive turn in the N-termini hairpin leads to a detectable intermediate for protein G

Running ~20 *μ*s long replica exchange simulations we found only 2 walkers out of the 30 eventually fold into the native structure (Fig. S2). We use the fraction of native contacts formed (*Q*)^34^ to analyze the evolution of the folding process, finding very similar folding patterns in both walkers (see Fig.2a and S4). We furthermore divide *Q* into each of the fragments relevant to the folding pathway: formation of the first hairpin (*Q*_12_), second hairpin (*Q*_34_), α-helix (*Q_α_*, and formation of the β-sheet (*Q*_1234_). Tracking the time evolution of these fragments, we identify the C-terminal hairpin first folds into its native form, followed by the appearance of a register shifted N-terminal hairpin alongside it (see Fig.2a). Formation of the two hairpins allows a native-like interaction between strands one and four, crucial for unfolding strand 2 and then refolding the N-terminal hairpin into the correct registry through the formation of the native turn. The helix, due to the local nature of the interaction, forms transient native like contacts along the folding trajectory. It is only after the four β strands are in place forming a β-sheet that the helix is stabilized by packing against it. The observed folding behavior from our simulations is in excellent agreement with observation from ψ analysis, in particular, the presence of non-native interactions in the N-terminal turn caused by the preference of a type I’ turn (N8-G9) over the native type I turn.

**Figure 2.**
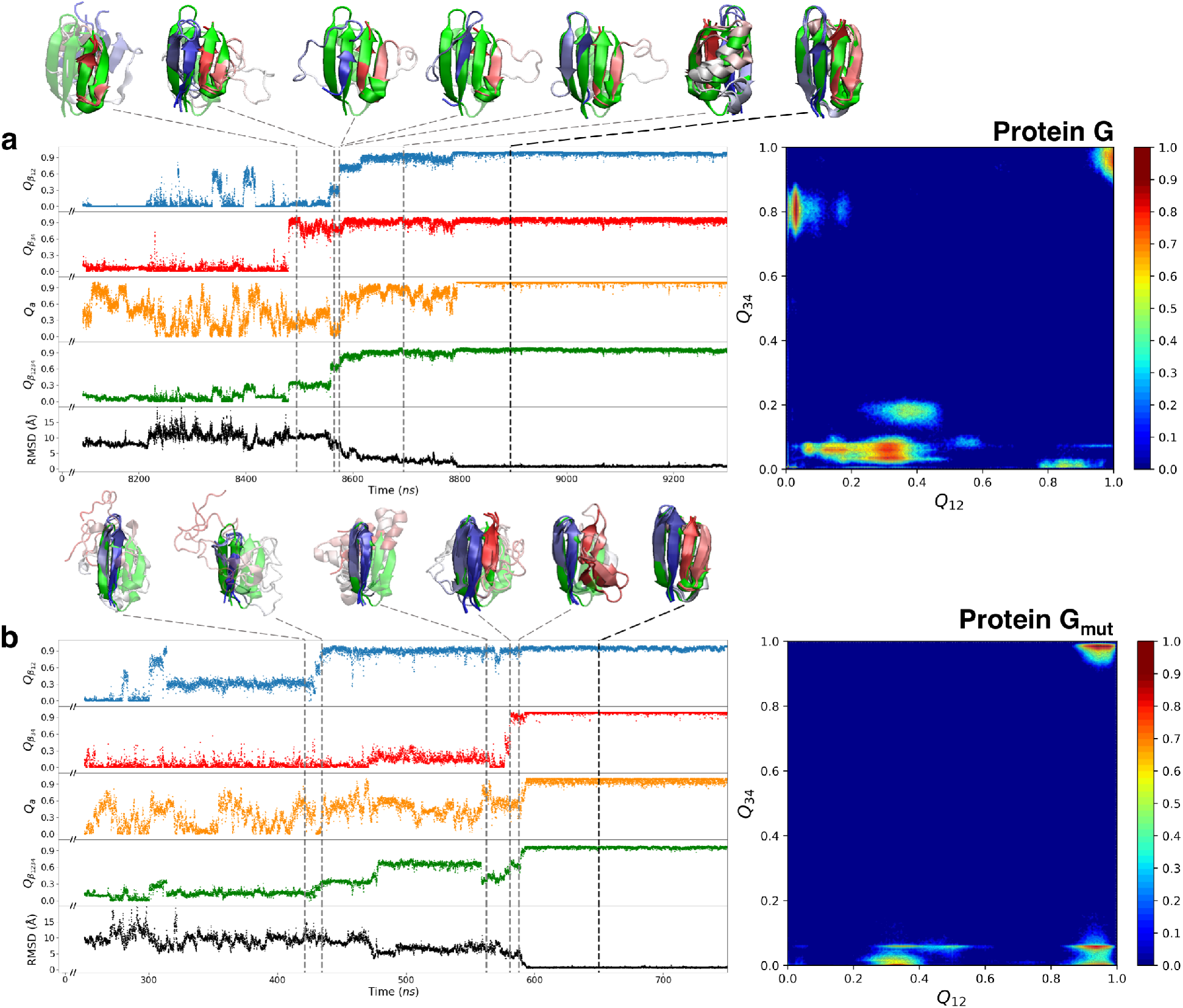
Fragments formation during folding of protein G and its mutant. **Left:** Native contact fractions for β_12_ (blue), β_34_ (red), α helix (orange), β_1234_ (green) and total RMSD (black) against native of protein G **(a)** and NuG2 **(b)** along the folding transition. Representative conformations are shown at different folding stages compared with native structure (green). **Right:** Normalized histogram of sampling at the lowest replica projected on the native contact fractions of N-terminal hairpin and C-terminal hairpin for protein G **(a)** and NuG2 **(b)**.

### Redesigning the first hairpin in NuG2 stabilizes the N-terminal hairpin and induces a change in the folding pathway with respect to protein G

In the designed protein G mutant (NuG2) there is no possibility for a type I turn. Thus, the N-terminal hairpin spontaneously forms through a type I’ turn (N8-G9), resulting in the major folding pathway going through the N-termini rather than the C-termini. Even with the correct turn, we identify a shifted registry hairpin formation before the native hairpin is formed. Although the C-terminal hairpin maintains the ability to fold independently, we find that interactions with the N-terminal hairpin accelerates the folding of the C-terminal hairpin through a near native interaction between the first and fourth β strands (Fig.2b). Only 1 out of the 30 walkers sampled the native state within ~20 *μ*s of simulation with high populations (Fig. S2). This folding behavior agrees well with recent analysis on explicit solvent MD trajectories on a triple mutant of NuG2b from the Shaw group^21,25,26^.

The Bayesian inference approach allows different replicas at different times to sample which of the possible regions of the protein favor extended conformations, while the nature of the flat-bottom restraints avoids biases when such extended conformations are satisfied. This allows us to analyze the relative populations of the hairpins seen in simulation. Analysis of the lowest temperature (300 K) replica (see the right panel in Fig.2b) shows that indeed the conformation with only the C-terminal hairpin was rarely found in the native state, while the N-termini can be observed in high populations in both the register-shifted and native forms. We observe the opposite trend for protein G (see right panel in Fig.2a). Folding of protein G proceeds through a major state in which the C-termini hairpin is present, but there are also large populations of the registry shifted N-termini hairpin where the C-termini hairpin is not formed. Such misfolded conformations are long-lived and prevent many walkers from reaching the native state (see Fig. S2). The bottleneck arises from the presence of a non-native turn that leads to the registry shifted conformation. Capturing this state is a feature of the current protocol, where biasing to individual strands allows any possible strand pairing beyond what we could predict initially. This state is reflective of force field preferences, and is compatible with known experimental data^7^.

### MELD acceleration is related to folding complexity

Protein L and L_mut_ are slower folders than protein G and Gmut. In our computational approach, they require shorter simulations to capture the native state, which was found in many independent walkers (13 and 18 walkers for protein L and its mutant respectively, see Fig. S3). Both proteins fold through the same N-terminal pathway and analysis of the low replica ensembles (see right panels in Fig.3) show the highest population states involve formation of the native structure and native N-terminal hairpin. For both proteins, there are few off-registry hairpin arrangements observed. Our results suggest that the design of the C-terminal hairpin is not likely to alter folding pathways, despite its faster folding rate^13^. As in the case of NuG2, interactions between the first and fourth β strands helped stabilize the formation of the second hairpin (C-terminal). (Fig.3) The slower experimental folding time of protein L/L_mut_ can be explained by a lower intrinsic stability of the strands. MELD simulations stabilize this critical step by imposing restraints on extended conformations, leading to more productive hairpin formation. Accelerating this initial nucleation step leads to a faster folding computational pipeline. Despite many folding events, we noticed the implicit solvent we used was not enough to fully stabilize the helix structure packed against the *β*-sheet, resulting in some of the RMSD distributions shifted to higher values than the threshold for native state (See Fig. S1a and S5).

**Figure 3.**
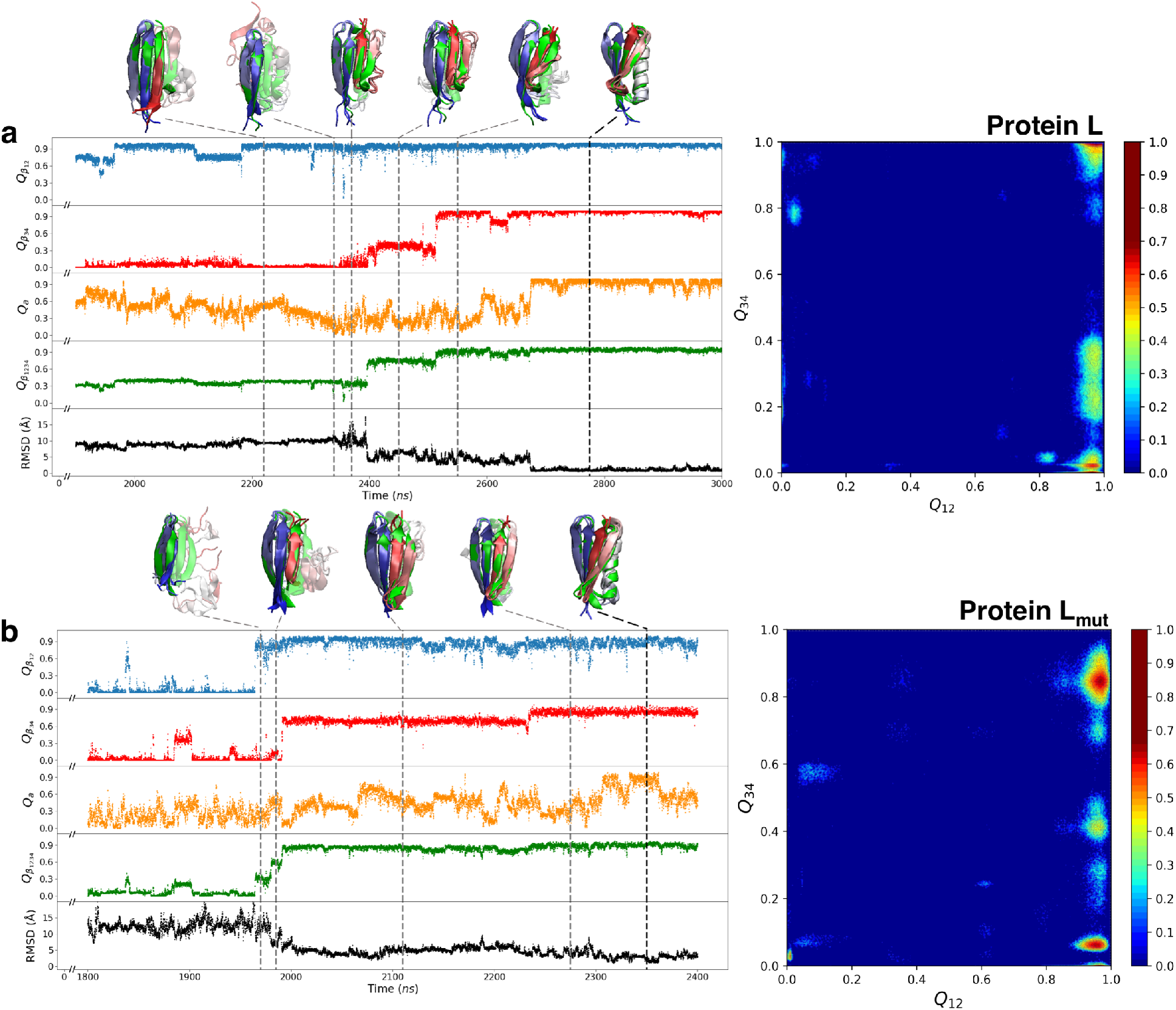
Fragments formation during folding of protein L and its mutant. **Left:** Native contact fractions for β_12_ (blue), β_34_ (red), α helix (orange), β_1234_ (green) and total RMSD (black) against native of protein L **(a)** and L mutant **(b)** along the folding transition. Representative conformations are shown at different folding stages compared with native structure (green). **Right:** Normalized histogram of sampling at the lowest replica projected on the native contact fractions of N-terminal hairpin and C-terminal hairpin for protein L **(a)** and L mutant **(b)**.

### Adaptive sampling combined with MSM analysis supports the folding mechanism for protein G and its mutant

MELD simulations provide an incomplete picture of the folding mechanism in protein G and its mutant due to few folding events. A more detailed picture emerges from aggregated milliseconds MD simulation generated by adaptive sampling with the same physical model (see methods) for protein G and its mutant. Using dimensionality reduction techniques based on time-lagged independent component analysis(tICA, see methods), a funnel-like folding energy landscape emerges for both proteins (Fig.4a). For protein G, the tendency for early formation of C-terminal hairpin is clearly indicated by clustered microstates in the unfolded state and near the folded state region. Along the first independent component, native interactions between the first and fourth strands are present early on and a registry shifted behavior on the N-terminal identified from MELD simulation is also seen here to slow the folding process (the second-slowest timescale in NuG2). The designed new turn creates an intermediate state for conformations containing the native N-terminal hairpin for NuG2 before folding into the native structure. In contrast with protein G, the formation of the C-terminal hairpin becomes the critical step for folding, which is likely hindered by interactions between the first and fourth strand in antiparallel orientation observed in MELD simulations (see SI Fig. S6). A recent study on the triple mutant of NuG2 found such conformation has a lower internal energy compared with the native structure, but higher energy when solvation effects are taken into account^36^.

**Figure 4.**
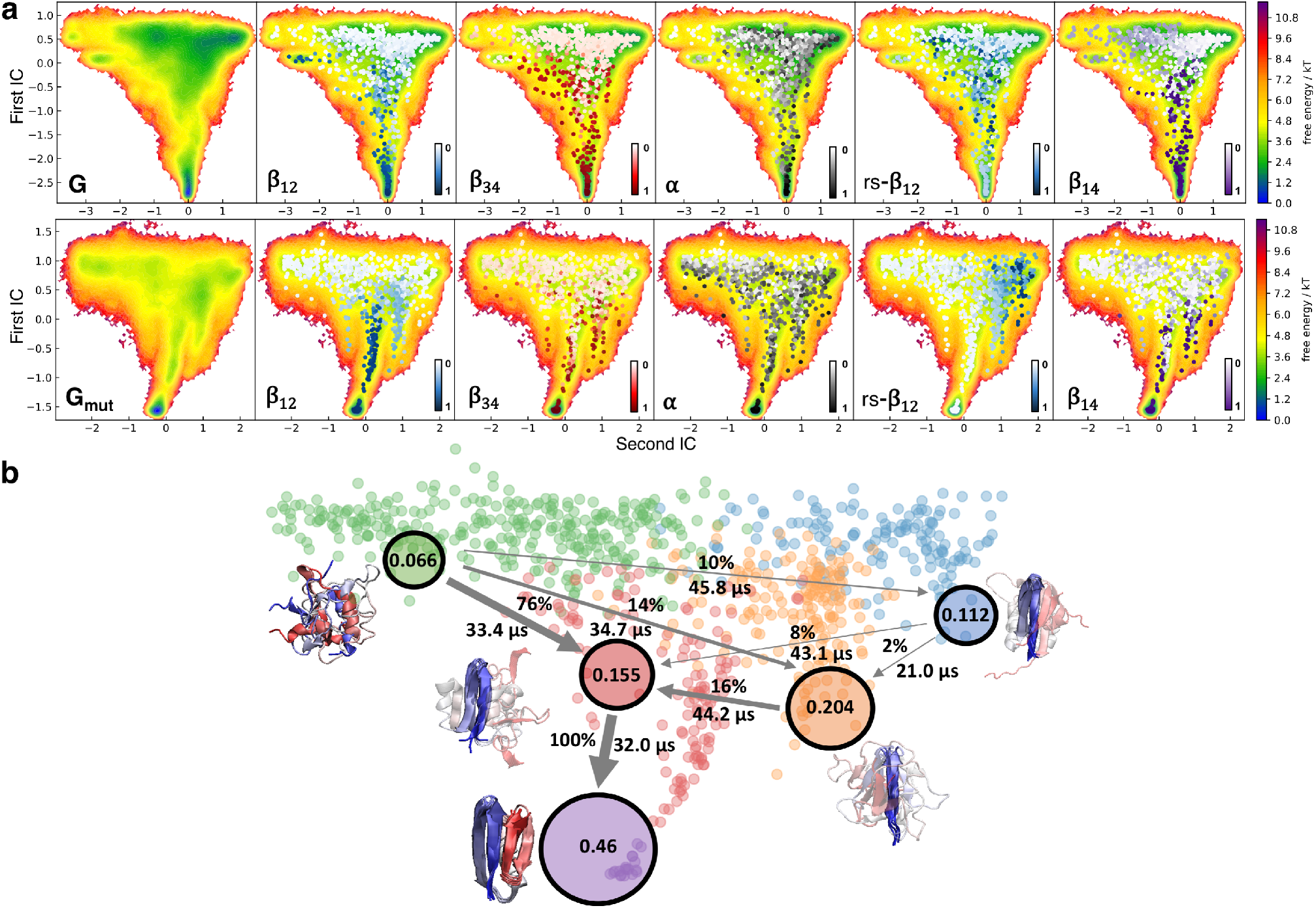
Folding energy landscape and kinetics from MSM analysis. **a** Two-dimensional representation of the free energy landscape projected on the first two independent components with native contact fraction of β_12_, β_34_, α helix, register shifted β_12_, and β_14_ shown on each clustered microstate for protein G (top) and G mutant (bottom). **b** Coarse-grained modeling on 800 clustered microstates with hidden markov model for protein G mutant. Flux network and mean first passage time are estimated from unfolded state (green) to folded state (purple) by transition path theory.

We coarse-grained the clustered microstates into five macro-states using a hidden Markov model (HMM) and applied transition path theory to determine the major folding pathways for NuG2. (see methods and Fig.4b) The initial state is defined on conformations with native contact fraction close to zero for all fragments, and the final state is the folded native state compared with the experimental structure. Unsurprisingly, the pathway with the largest flux went from the initial unfolded state to the state containing the native N-terminal hairpin, followed by the formation of the C-terminal hairpin and α-helix. Other pathways proceeded through conformations with the register shifted effect first before folding into native state. Figure S7 summarizes the Chapman-Kolmogorov results to pick a lagtime for the MSM as well as the mean first passage time between states. The pathway previously reported to form through the C-terminal hairpin first is not observed here^26^. A similar analysis for protein G (see Fig. S8) identifies 2 major states: folded and unfolded. As expected, the transition between the two states is significantly longer (hundreds of microseconds) than the one observed for the NuG2 mutant. While these results agree well with the experimental folding time, additional adaptive sampling is required for converged analysis of protein G (beyond the scope of the current work).

### AlphaFold predictions identify bottlenecks along the folding pathways

AlphaFold’s success in structure prediction (protein, protein-protein, and peptide-protein) poses multiple questions about how much of the biophysics encoded through machine learning. While AF has not been designed to capture protein folding pathways, we wondered if through the training process AF is capable of bringing insights to protein folding pathways by characterizing the structure of different building blocks that lead to the folded state. We start with the knowledge that there are four β-strands and one α-helix and ask AF to make predictions for different combinations of connected secondary structure elements in the native structure (Fig.5). We first look at predictions for the two hairpin intermediates and the α-helix for each sequence. Surprisingly, the N-termini hairpin of protein G as well as the C-termini hairpin of protein L are correctly predicted to be non-native (Fig.5). In the first case, AF reflects the preference for the type I turn leading to a non-native hairpin, whereas the designed type I’ turn is predicted in the native form of NuG2. In addition, using strands 1, 2 and 4 for prediction identifies the possible antiparallel formation between strand 1 and 4 that appears to be the dominant conformation in the intermediate state found in our MD simulation (see Figs. S6 and S9).

**Figure 5.**
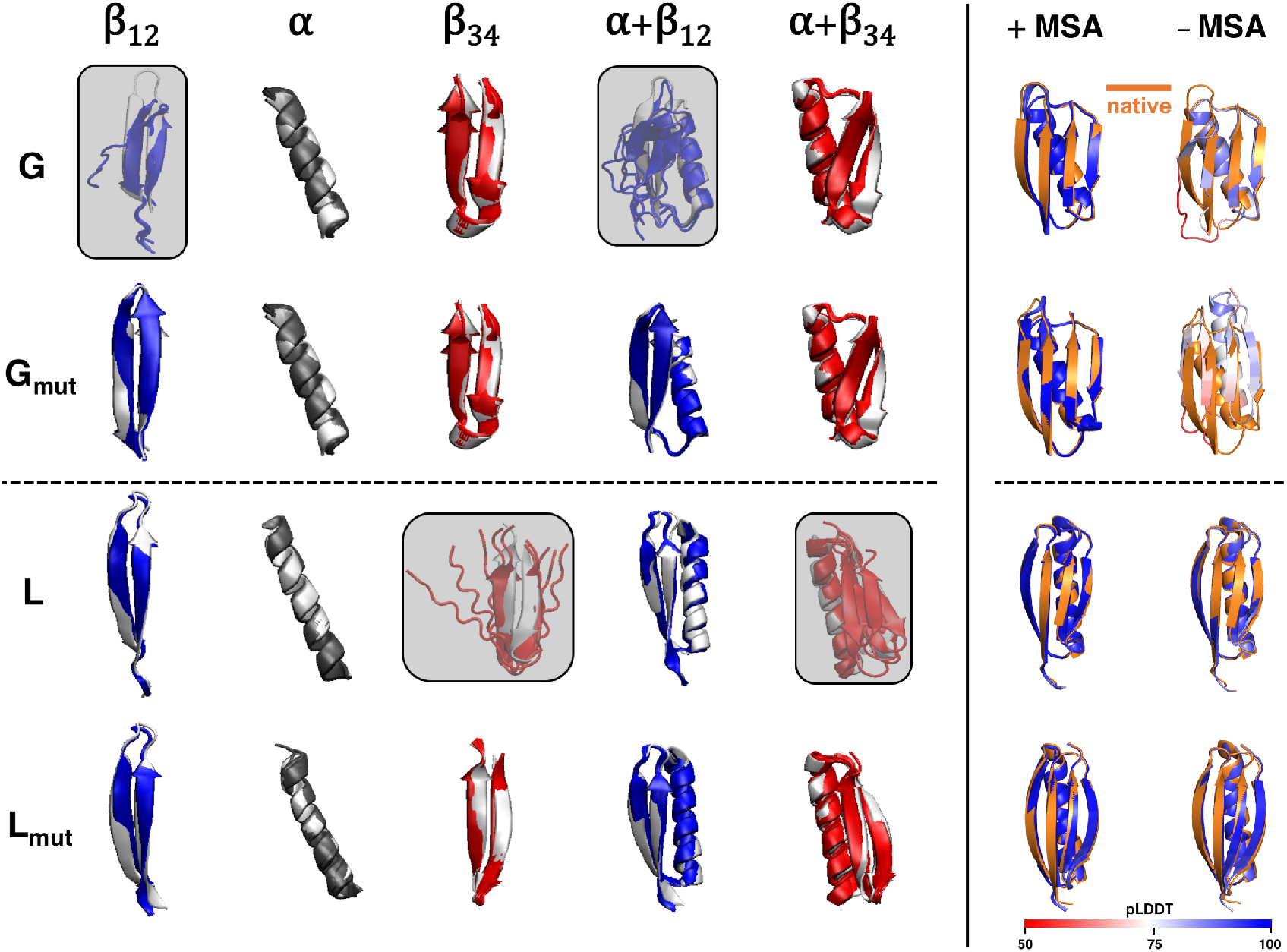
Protein fragment and full structure predictions from AlphaFold. **Left:** Structures of five AF predicted models for different fragments (β_12_ (blue), α helix (dark grey), β_34_ (red), β_12_ with α helix, and α helix with β_34_) are aligned to the native structure (white). **Right:** Full structure predictions colored by pLDDT score aligned with the native structure (orange) with and without MSA as input. (Only the top model ranked by pLDDT score is shown here for prediction without MSA, see Fig. S10 for all five models.)

In the case of protein L, the C-terminal hairpin is predicted to be unstructured, whereas the mutant with the redesigned C-terminal is predicted to be a hairpin. For all proteins, the sequence of the helix is predicted to be helical even by itself, contrary to our observations in MD simulations or the experimental TSE analysis. This is likely due to over-represented helical structures in the training dataset, which have also been observed in the recent benchmark on predicting protein-peptide structures with AF^39^. The inclusion of MSA as input does not affect local secondary structure prediction (e.g. β_12_, see Fig.5 and S10), but the accuracy for prediction of long-range interactions between secondary structure fragments dropped without coevolution information (e.g. β_12_ and α-helix). A recent study suggests that coevolution data helps solve the global search problem by finding a low energy conformation that is then optimized along the learned biophysical function^40^. This agrees well with our observations, where AF encodes the structural pattern for local interactions, while the MSA governs the accuracy of predictions for long-range effect (residues far along the sequence that are in contact). The same conclusions are observed from predictions for the full sequence using coevolution information (+MSA) or only the protein sequence (–MSA in Fig.5). While all five AF models are predicted almost identical to experiments using MSA, only the local structure preference is captured using a single query sequence. As in the case of protein G, the N-terminal hairpin is predicted to contain a non-native type I’ turn in the top-ranked model and reamins unstructured in other models (see Fig.5 and S10). Interestingly, the predicted disparity between G/G_mut_ and L/L_mut_ without MSA is in agreement with the folding complexity difference found from our MD simulation study. Together with other recent studies, these results increase the interpretability of the trained neural network AF represents. In particular, the current results show a finely tuned sensitivity to the folding events observed using MD simulation.

## Discussion

The formation of intra-strand atomic interactions together with the hydrophobic effect favor the folding process, while chain entropy favors unfolded ensembles^23^. Along the folding process, intra-strand interactions are sampled with a certain probability (e.g. closer native contacts have a higher probability) and once established have a certain lifetime depending on the surrounding environment. When enough such events have accumulated, the protein reaches the transition state, sampling the native state soon after. By their nature, molecular dynamics simulations can capture this process, the question is whether they lead to a similar TSE and intermediates on its way to the folded state. The challenge are the simulation timescales of these folding processes, which range from the microsecond to second timescale for the four proteins studied here. For the fastest folding protein in the set (NuG2) there are two sets of conventional MD starting from unfolded simulations for its triple mutant form. While in one study, running at the melting temperature the native state is found^25^, simulations in implicit solvent in the 50 *μ*s range did not sample the correct topology with T-REMD^41^. On the other hand, extensive sampling (50 ms aggregated sampling time) using distributed computing coupled with MSM analysis was able to identify possible folding pathways for the slower folding protein G^8^.

Here we have shown that a unified protocol based on evidence from experimental ψ value analysis is enough to distinguish the folding pathways of protein G and L as well as their mutants. We believe this approach offers advantages over current methodologies to connect experimental information on pathways with atomistic detail. We follow the philosophy of the smallest perturbation to the system, thus we guide towards the formation of extended conformations for core residues belonging to strands. In this way, interactions between different strands can lead to productive β-strand pair formation and ultimately help nucleate global folding. The method is sensitive to the nuances in protein pathways from there on, and we expect that complex folding pathways will still require long simulation time.

By biasing only a few residues to extended regions in the Ramachandran plot we impose no information regarding how the strands should pair (local or global interactions), their registry or even their parallel/antiparallel orientation. In this way, the pathways found by MELD are not biased. The folding complexity leads to different acceleration rates inside MELD. Protein L/L_mut_ show little frustration after the first hairpin is formed, rapidly leading to the TSE and global fold. The slower acceleration for protein G and its mutant reflects the higher complexity in the energy landscape that proceeds through intermediate states. Our results identify the correct pathway in all four cases. Hence, the physics model coupled with the Bayesian inference based sampling strategy are sensitive to sequence preferences along the folding mechanism.

We observed a competitive turn in the N-terminal hairpin of protein G playing an important role during folding. Its formation leads to an interaction between strands 1 and 4 which is native like. Eventually, strand 2 refolds into the correct hairpin and the native β-sheet is established. Finally, the alpha-helix is stabilized by packing against the β-sheet. The local nature of α-helix formation is seen through frequent transient native interactions that are only stabilized once the helix packs against the four strands. The non-native behavior caused by the competitive turn was previously detected in the ψ analysis for the TSE^7^. The engineering of protein G to its mutant stabilizes the first hairpin, which indeed now nucleates the folding of the protein. In a twist, the second hairpin is now able to pack competitively against the first hairpin leading to an intermediate. Thus, despite the fastest folding time in the dataset, the folding pathway is not purely two state as there is a detectable intermediate before the TSE is formed. Unlike protein G and its variant, there are no detectable intermediates with competitive turns or a hairpin registry shift to slow down folding for protein L and its mutant. However, the observed high population for conformations with an N-terminal hairpin could be viewed as an on-path intermediate that supports the recent observation by single-molecule fluorescence resonance energy transfer (FRET) spectroscopy^17^.

Deep neural network based models forgo the need for expensive sampling by training on sets of known protein structures. Despite their unprecedented impact, interpretability of what the network has learned, its limits and possibilities are a matter of active research in the field. A recent work proposes that MSA and the encoded energy function help AF identify regions of conformational space that are protein-like – whereas the network has learned a sort of biophysical energy function that allows it to drive the structure towards the global minima^40^. Interestingly, breaking the protein into short fragments and asking AF for structural models reveals sensitivity to the type of turn (type I or type I’). While more analysis is needed, tools such as AF might provide information about possible structured fragments that lead to intermediates as working hypothesis in the absence of experimental data for folding pathways. A minimal set of restraints could in turn be used within MELD as outlined in this work to characterize possible folding mechanism.

## Methods

### MELD folding simulations

MELD is a flexible Bayesian framework for accelerating physics-based molecular simulation guided by external information. Under the formulation of Bayes’ theorem, *p*(*x* | *D*)~*p*(*D* | *x*)*p*(*x*), the prior probability of a conformation *x*, *p*(*x*), is the Boltzmann probability distribution determined by the selected force field. The folding simulation is accelerated given a map of the four native β strands as guidance. The likelihood of the such guidance *D* given *x*, *p*(*D* | *x*), is proportional to *e*^−*E_r_*(*x*)/*kT*^, where *T* is the temperature and *E_r_*(*x*) is the secondary structure restraint energy from MELD. In addition, given each sampled conformation *x*, the restraint energy is ranked for all four β strands, and only 50% of the restraints (those with lowest energy values) are activated. The restraint energy is defined as a flat-bottom harmonic potential with zero restraint energy in the region compatible with guidance, thus the relative probability of forming N-terminal hairpin and C-terminal hairpin is consistent with unbiased simulations because both require the conformation to match 50% of all guidance. The Hamiltonian and temperature replica exchange MD (H,T-REMD) is the sampling engine in MELD to further avoid kinetic traps. We keep the same force constant for all replicas and set the temperature range from 300 to 450 K with geometric temperature scaling. All simulations were done with OpenMM^42^ started from fully extended chains in all atomic detail parameterized by Amber ff14SB sidechain^41,43^ + ff99SB backbone^44^ with the implicit solvent model GB-neck2^45^. The hydrogen mass repartitioning^46^ was used to allow 4.5-fs timestep.

### MELD folding restraints

To nucleate the formation of the strands we used the information that there are four possible strands in each protein. The ones in protein L (and mutant) are longer than the ones in protein G (and mutant). Inducing a strand like conformation requires the dihedrals of different residues to be in the β region of the Ramachandran plot as well as some correlation between their values. We induce this correlation by combining flat bottom harmonic restraints to the β region and distance restraints (1-4 and 1-5) to distances compatible with β strands. What is important is that for each simulation the 4 strands have equal number of possible restraints (equal nucleating length). We choose two overlapping fragments for protein G and is mutant in each of its strand and three overlapping fragments for protein L and its mutant (see setup file in https://github.com/PDNALab/GL_folding).

### Adaptive sampling simulations

The first round of adaptive sampling for both protein G and its mutant was initialized with a batch of 200 ns simulations starting from seeds selected among high temperature simulations (unfolded state) and the native structure (folded state). Then new seeds were spawned for the next round from microstates with low counts in the transition matrix from a MSM estimated based on sampling from the previous round. In total, we generated ~1.6 ms simulations for NuG2 and ~3 ms for protein G as its simulations were extended to 400 ns to obtain higher statistical significance based on its slower folding time. All adaptive sampling used the pmemd.cuda engine^47^ in AMBER^48^ at 300 K under the same force field and solvent model described for MELD simulations. The hydrogen mass repartitioning was used to allow 4-fs timestep, with trajectories written every 1 ns for analysis.

### MSM analysis and coarse-grain modeling

The adaptive sampling datasets for protein G and its mutant are analyzed with the PyEMMA software^49^ for each system. The pairwise inter-residue distances, RMSD against the native state, and the native contact fractions for protein fragments were included as input features. Then, the seven slowest components were selected from dimension reduction on those features using time-lagged independent component analysis (tICA) with 50 ns as lag time and k-means clustering was employed to obtain 800 microstates. The validation of the Markov model on those microstates were performed by a Chapman–Kolmogorov test on the final model with the selected lag time (Fig. S7a).

### AlphaFold predictions

Fragment predictions were performed using a local installation of the publicly available ColabFold^50^, with trained parameters from five models of AF. No template or additional refinement were used for the predictions. For disconnected fragments such as β−124, we used a poly-glycine linker, an idea that has already been used for modeling protein-peptide complex using AF^39^, to connect β−12 and β−4 for monomer prediction.

## Acknowledgements

We thank Ken A. Dill for critical reading and helpful suggestions. We are thankful to computational resources from XSEDE (TG-BIO210099) and the HiPerGator supercomputer at the University of Florida.

## Author contributions statement

L.C and A.P conceived and planned the research, L.C. conducted and analyzed the simulations, L.C. and A.P. discussed the results, drafted and reviewed the manuscript.

## References

1. Jumper, J. et al. Highly accurate protein structure prediction with AlphaFold. Nature 1–11 (2021).

2. Baek, M. et al. Accurate prediction of protein structures and interactions using a three-track neural network. Science eabj8754 (2021).

3. Tunyasuvunakool, K. et al. Highly accurate protein structure prediction for the human proteome. Nature 1–9 (2021).

4. Anishchenko, I. et al. De novo protein design by deep network hallucination. Nature 1–6 (2021).

5. Levinthal, C. Are there pathways for protein folding? J. de chimie physique 65, 44–45 (1968).

6. McCallister, E. L., Alm, E. & Baker, D. Critical role of -hairpin formation in protein G folding. Nat. Struct. Biol. 7, 669–673 (2000).

7. Baxa, M. C. et al. Even with nonnative interactions, the updated folding transition states of the homologs Proteins G & L are extensive and similar. Proc. Natl. Acad. Sci. 112, 8302–8307 (2015).

8. Lapidus, L. et al. Complex Pathways in Folding of Protein G Explored by Simulation and Experiment. Biophys. J. 107, 947–955 (2014).

9. Kim, D. E., Fisher, C. & Baker, D. A Breakdown of Symmetry in the Folding TransitionState of Protein L.pdf. J. Mol. Biol. 298, 971–984 (2000).

10. Yoo, T. Y. et al. The Folding Transition State of Protein L Is Extensive with Nonnative Interactions (and Not Small and Polarized). J. Mol. Biol. 420, 220–234 (2012).

11. Maity, H. & Reddy, G. Folding of Protein L with Implications for Collapse in the Denatured State Ensemble. J. Am. Chem. Soc. 138, 2609–2616 (2016).

12. Maity, H. & Reddy, G. Transient intermediates are populated in the folding pathways of single-domain two-state folding protein L. J. Chem. Phys. 148, 165101 (2018).

13. Kuhlman, B., O’Neill, J. W., Kim, D. E., Zhang, K. Y. & Baker, D. Accurate computer-based design of a new backbone conformation in the second turn of protein L. J. Mol. Biol. 315, 471–477 (2002).

14. Sheinerman, F. B. & Brooks, C. L. Calculations on folding of segment B1 of streptococcal protein G. J. Mol. Biol. 278, 439–456 (1998).

15. Karanicolas, J. & Brooks, C. L. The origins of asymmetry in the folding transition states of protein L and protein G. Protein Sci. 11, 2351–2361 (2002).

16. Morrone, A. et al. GB1 Is Not a Two-State Folder: Identification and Characterization of an On-Pathway Intermediate. Biophys. J. 101, 2053–2060 (2011).

17. Aviram, H. Y., Pirchi, M., Barak, Y., Riven, I. & Haran, G. Two states or not two states: Single-molecule folding studies of protein L. J. Chem. Phys. 148, 123303 (2017).

18. Brown, S. & Head-Gordon, T. Intermediates and the folding of proteins L and G. Protein Sci. 13, 958–970 (2004).

19. Cheng, Q., Joung, I., Lee, J., Kuwajima, K. & Lee, J. Exploring the Folding Mechanism of Small Proteins GB1 and LB1. J. Chem. Theory Comput. 15, 3432–3449 (2019).

20. Bitran, A., Jacobs, W. M. & Shakhnovich, E. Validation of DBFOLD: An efficient algorithm for computing folding pathways of complex proteins. PLoS Comput. Biol. 16, e1008323 (2020).

21. Mitsutake, A. & Takano, H. Folding pathways of NuG2—a designed mutant of protein G—using relaxation mode analysis. J. Chem. Phys. 151, 044117 (2019).

22. Adhikari, U. et al. Computational Estimation of Microsecond to Second Atomistic Folding Times. J. Am. Chem. Soc. 141, 6519–6526 (2019).

23. Nauli, S., Kuhlman, B. & Baker, D. Computer-based redesign of a protein folding pathway. Nat. Struct. Biol. 8, 602–605 (2001).

24. Nauli, S. et al. Crystal structures and increased stabilization of the protein G variants with switched folding pathways NuG1 and NuG2. Protein Sci. 11, 2924–2931 (2002).

25. Lindorff-Larsen, K., Piana, S., Dror, R. O. & Shaw, D. E. How Fast-Folding Proteins Fold. Science 334, 517–520 (2011).

26. Schwantes, C., Shukla, D. & Pande, V. Markov State Models and tICA Reveal a Nonnative Folding Nucleus in Simulations of NuG2. Biophys. J. 110, 1716–1719 (2016).

27. Sugita, Y. & Okamoto, Y. Replica-exchange molecular dynamics method for protein folding. Chem. Phys. Lett. 314, 141–151 (1999).

28. Hruska, E., Abella, J. R., Nüske, F., Kavraki, L. E. & Clementi, C. Quantitative comparison of adaptive sampling methods for protein dynamics. J. Chem. Phys. 149, 244119 (2018).

29. Zuckerman, D. M. & Chong, L. T. Weighted Ensemble Simulation: Review of Methodology, Applications, and Software. Annu. Rev. Biophys. 46, 1–15 (2016).

30. Bussi, G. & Laio, A. Using metadynamics to explore complex free-energy landscapes. Nat. Rev. Phys. 2, 200–212 (2020).

31. Scalley, M. L. et al. Kinetics of Folding of the IgG Binding Domain of Peptostreptoccocal Protein L. Biochemistry 36, 3373–3382 (1997).

32. MacCallum, J. L., Perez, A. & Dill, K. A. Determining protein structures by combining semireliable data with atomistic physical models by Bayesian inference. Proc. Natl. Acad. Sci. 112, 6985–6990 (2015).

33. Perez, A., MacCallum, J. L. & Dill, K. A. Accelerating molecular simulations of proteins using Bayesian inference on weak information. Proc. Natl. Acad. Sci. 112, 11846–11851 (2015).

34. Best, R. B., Hummer, G. & Eaton, W. A. Native contacts determine protein folding mechanisms in atomistic simulations. Proc. Natl. Acad. Sci. 110, 17874–17879 (2013).

35. Pérez-Hernández, G., Paul, F., Giorgino, T., Fabritiis, G. D. & Noé, F. Identification of slow molecular order parameters for Markov model construction. J. Chem. Phys. 139, 015102 (2013).

36. Maruyama, Y. & Mitsutake, A. Stability of Unfolded and Folded Protein Structures Using a 3D-RISM with the RMDFT. J. Phys. Chem. B 121, 9881–9885 (2017).

37. Noé, F., Wu, H., Prinz, J.-H. & Plattner, N. Projected and hidden Markov models for calculating kinetics and metastable states of complex molecules. J. Chem. Phys. 139, 184114 (2013).

38. Noé, F., Schütte, C., Vanden-Eijnden, E., Reich, L. & Weikl, T. R. Constructing the equilibrium ensemble of folding pathways from short off-equilibrium simulations. Proc. Natl. Acad. Sci. 106, 19011–19016 (2009).

39. Tsaban, T. et al. Harnessing protein folding neural networks for peptide–protein docking. Nat. Commun. 13, 176 (2022).

40. Roney, J. P. & Ovchinnikov, S. State-of-the-Art Estimation of Protein Model Accuracy using AlphaFold. bioRxiv 2022.03.11.484043 (2022).

41. Nguyen, H., Maier, J., Huang, H., Perrone, V. & Simmerling, C. Folding Simulations for Proteins with Diverse Topologies Are Accessible in Days with a Physics-Based Force Field and Implicit Solvent. J. Am. Chem. Soc. 136, 13959–13962 (2014).

42. Eastman, P. et al. OpenMM 7: Rapid development of high performance algorithms for molecular dynamics. PLoS Comput. Biol. 13, e1005659 (2017).

43. Maier, J. A. et al. ff14SB: Improving the Accuracy of Protein Side Chain and Backbone Parameters from ff99SB. J. Chem. Theory Comput. 11, 3696 – 3713 (2015).

44. Hornak, V. et al. Comparison of multiple Amber force fields and development of improved protein backbone parameters. Proteins 65, 712 – 725 (2006).

45. Nguyen, H., Roe, D. R. & Simmerling, C. Improved Generalized Born Solvent Model Parameters for Protein Simulations. J. Chem. Theory Comput. 9, 2020 – 2034 (2013).

46. Hopkins, C. W., Grand, S. L., Walker, R. C. & Roitberg, A. E. Long-Time-Step Molecular Dynamics through Hydrogen Mass Repartitioning. J. Chem. Theory Comput. 11, 1864–1874 (2015).

47. Gotz, A. W. et al. Routine Microsecond Molecular Dynamics Simulations with AMBER on GPUs. 1. Generalized Born. J. Chem. Theory Comput. 8, 1542–1555 (2012).

48. Case, D. A. et al. AMBER 2020. Univ. California, San Francisco (2020).

49. Scherer, M. K. et al. PyEMMA 2: A Software Package for Estimation, Validation, and Analysis of Markov Models. J. Chem. Theory Comput. 11, 5525–5542 (2015).

50. Mirdita, M. et al. ColabFold - Making protein folding accessible to all. bioRxiv 2021.08.15.456425 (2022).

