## Supplemental figures for "A consensus view on the folding mechanism of protein G, L and their mutants"

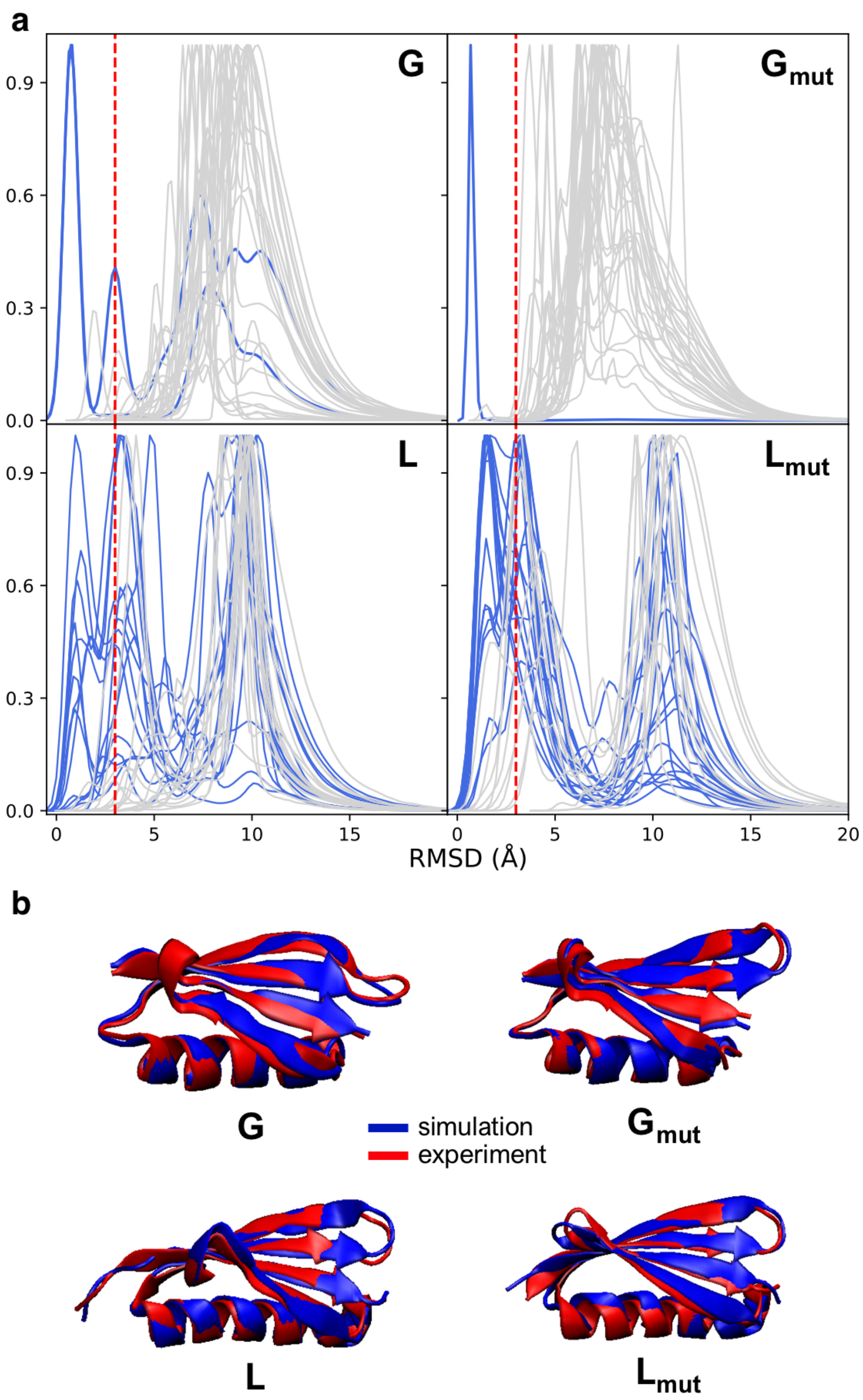


**Supplementary Figure 1. Summary of all "walkers" and representative structures in MELD simulations.** **a** Normalized histogram of RMSD against native structure for the 30 "walkers" in MELD simulations. Blue line indicates the walkers with high population in the native state (3 Å red dotted line). **b** Representative structures from MELD simulations by using hierarchical clustering at the lowest replica with Cpptraj[1] and clustering protocol can be found here[2].

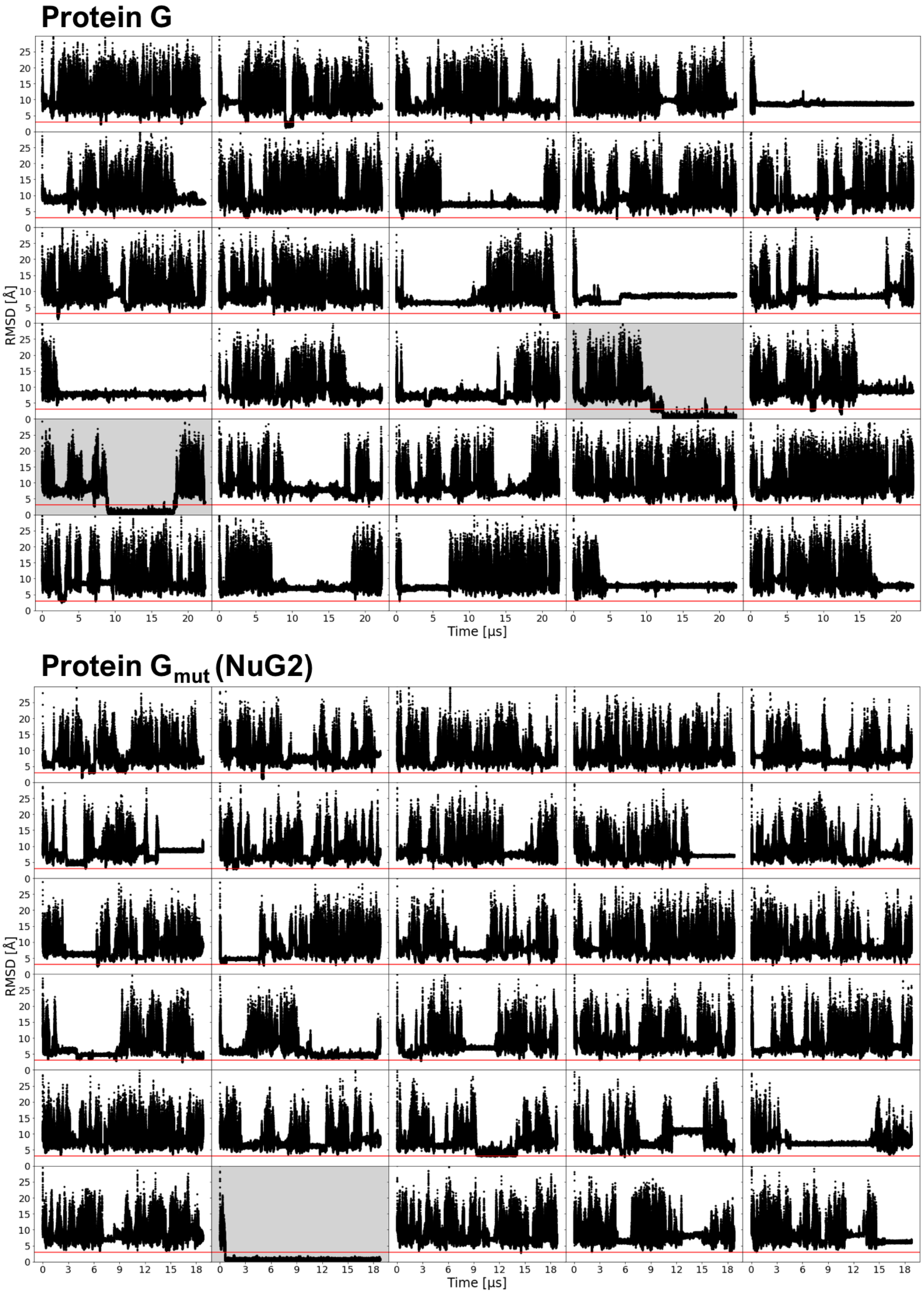


**Supplementary Figure 2. MELD folding simulation summary for G/G$_{mut}$** The time evolution for the RMSD of all walkers in the replica exchange simulation of protein G (top) and protein G mutant (bottom). Red line indicates the RMSD of 3 Å.

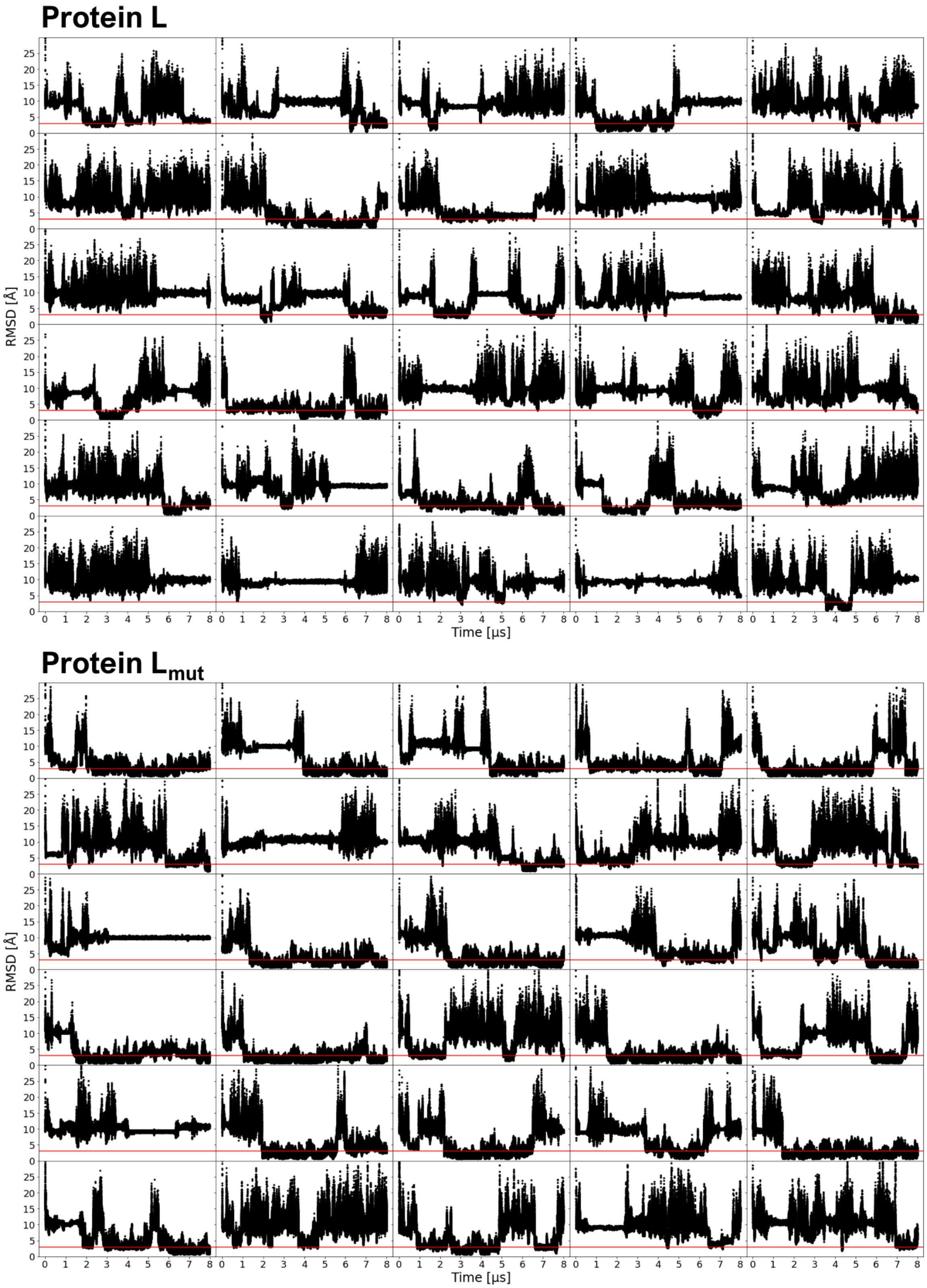


**Supplementary Figure 3. MELD folding simulation summary for L/$L_{mut}$** The time evolution for the RMSD of all walkers in the replica exchange simulation of protein L (top) and protein L mutant (bottom). Red line indicates the RMSD of 3 Å.

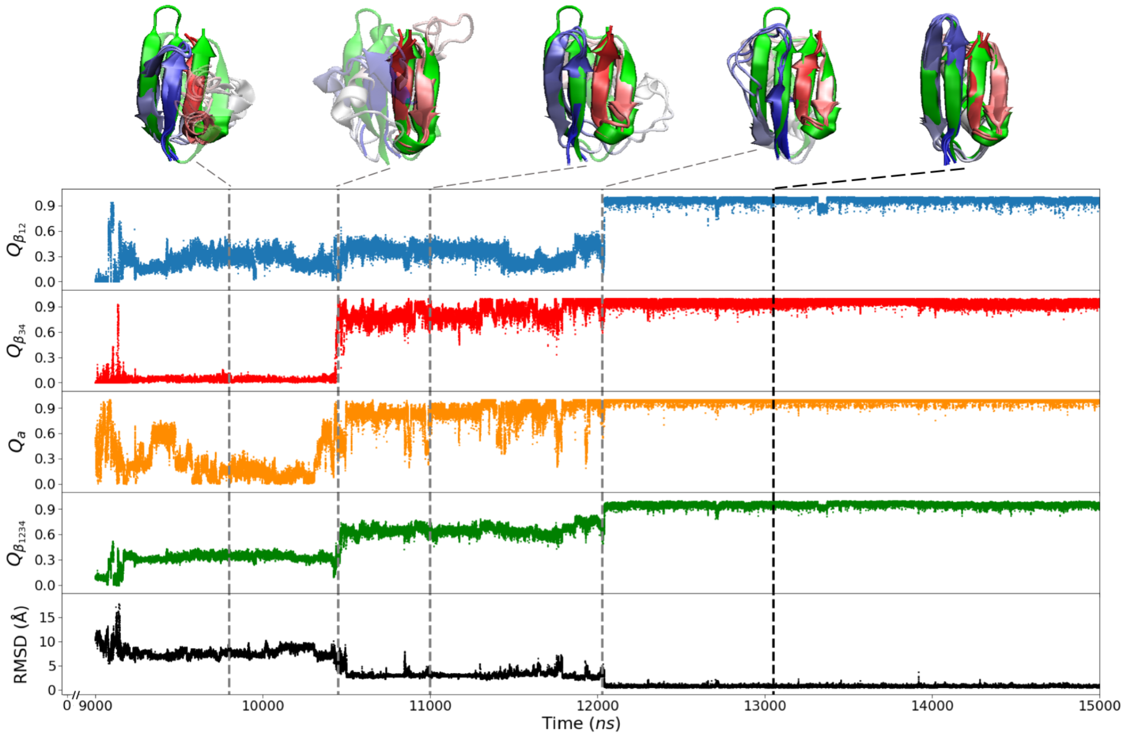


**Supplementary Figure 4. Transition from unfolded to folded state for protein G.** Native contact fractions for $\beta_{12}$ (blue), $\beta_{34}$ (red), $\alpha$ helix (orange), $\beta_{1234}$ (green) and total RMSD (black) for protein G simulation at replica 21. Representative conformations are shown at different stages along transition.

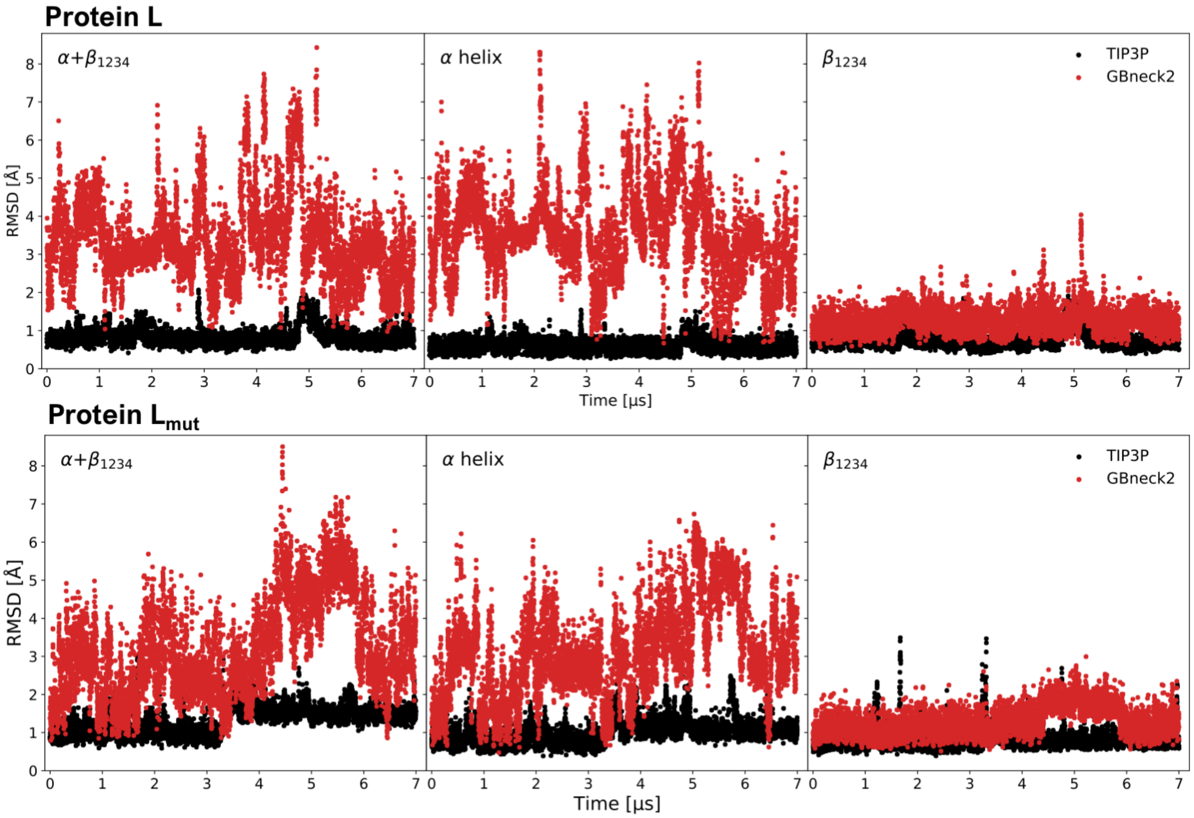


**Supplementary Figure 5. Instability of $\alpha$ helix in protein L/$L_{mut}$ in implicit solvent.** The time evolution of RMSD for full protein (left), $\alpha$ helix (middle) and $\beta_{1234}$ helix (bottom) simulated in explicit (black) and implicit solvent (red) at 300 K.

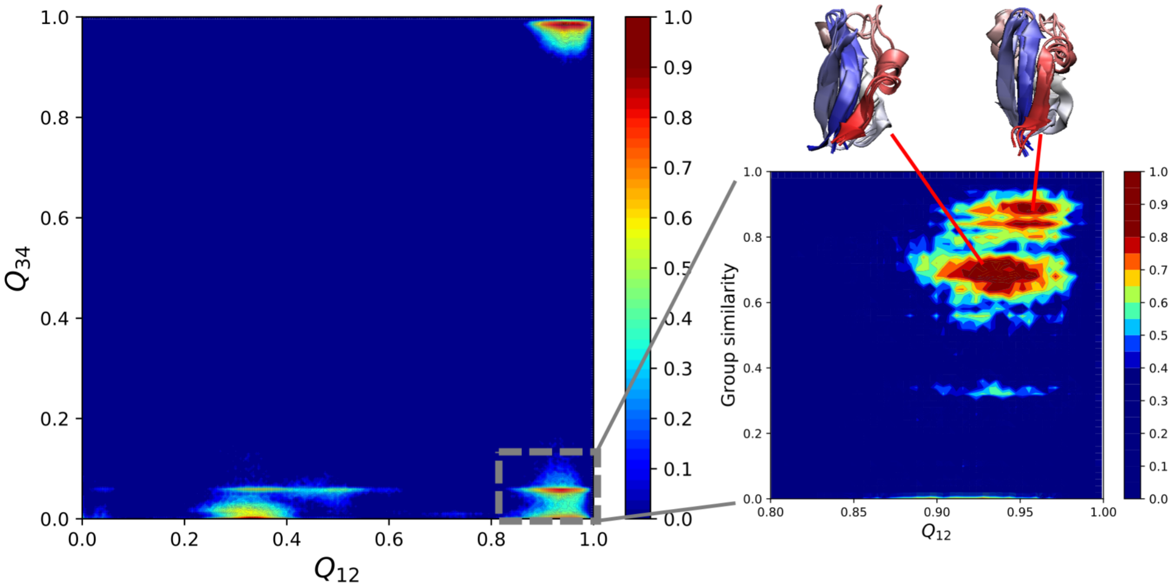


**Supplementary Figure 6. Dominant conformations of folding intermediates for NuG2.** Left: sampling at the lowest replica of simulation projected on the native contact fraction of the N-terminal and C-terminal hairpin, same with Fig. 2(b). Right: Dominant conformations represented by group similarity[3] based on contacts between $\beta_{12}$ and $\beta_4$ for sampled conformations with native contact fraction of the N-terminal hairpin.

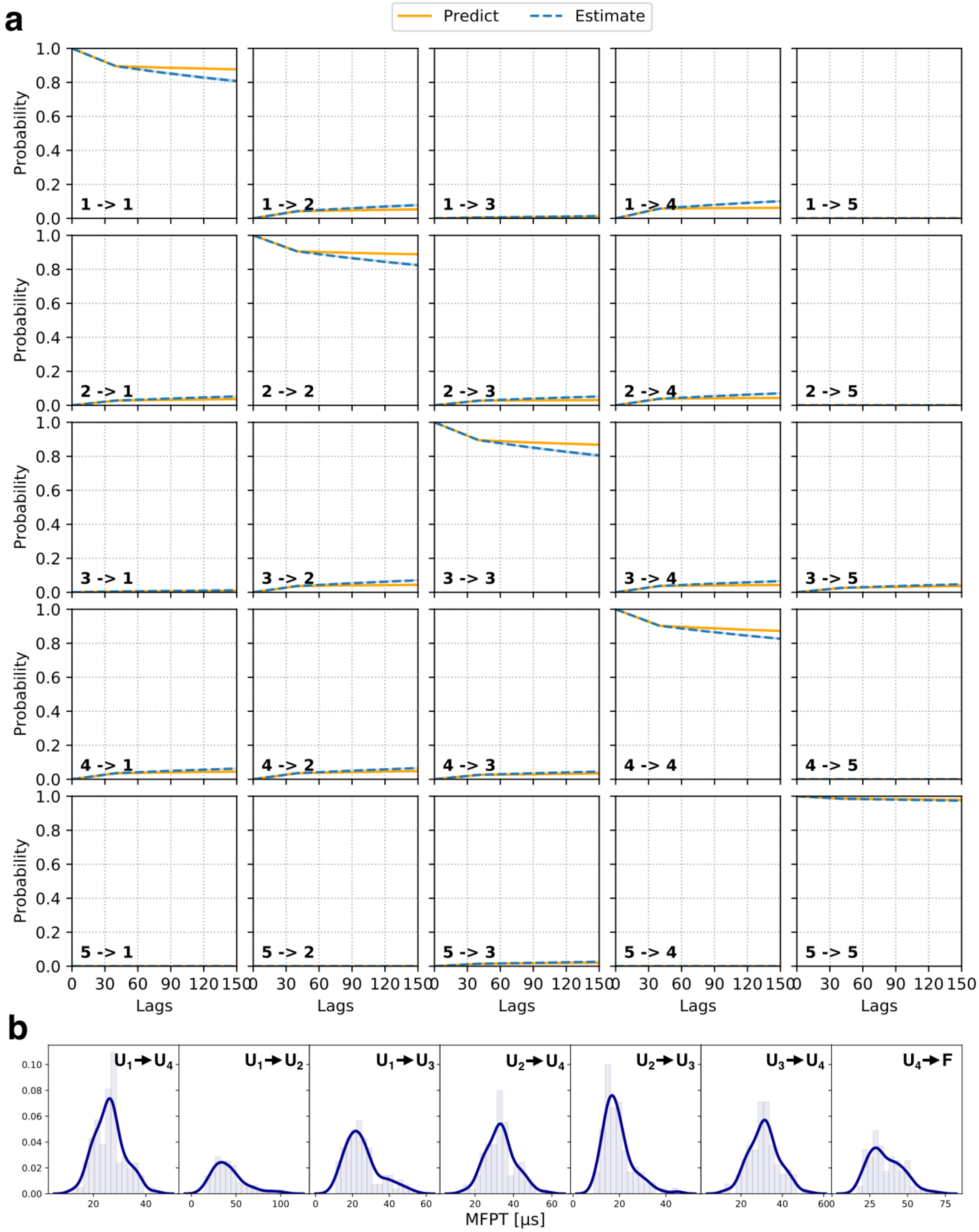


**Supplementary Figure 7. Validation and uncertainty of MSM analysis for protein $G_{mut}$.** **a** Chapman-Kolmogorov test for the final model of protein G mutant. Comparison of the transition probability for every pair of macrostates between the estimate (using MSM computed at various lag times) and the prediction from final model. **b** Sampled mean first passage time (MFPT) from the final model for transitions between macrostates ($U_1$ for green, $U_2$ for blue, $U_3$ for orange, $U_4$ for red and $U_5$ for purple) shown in Fig. 4b.

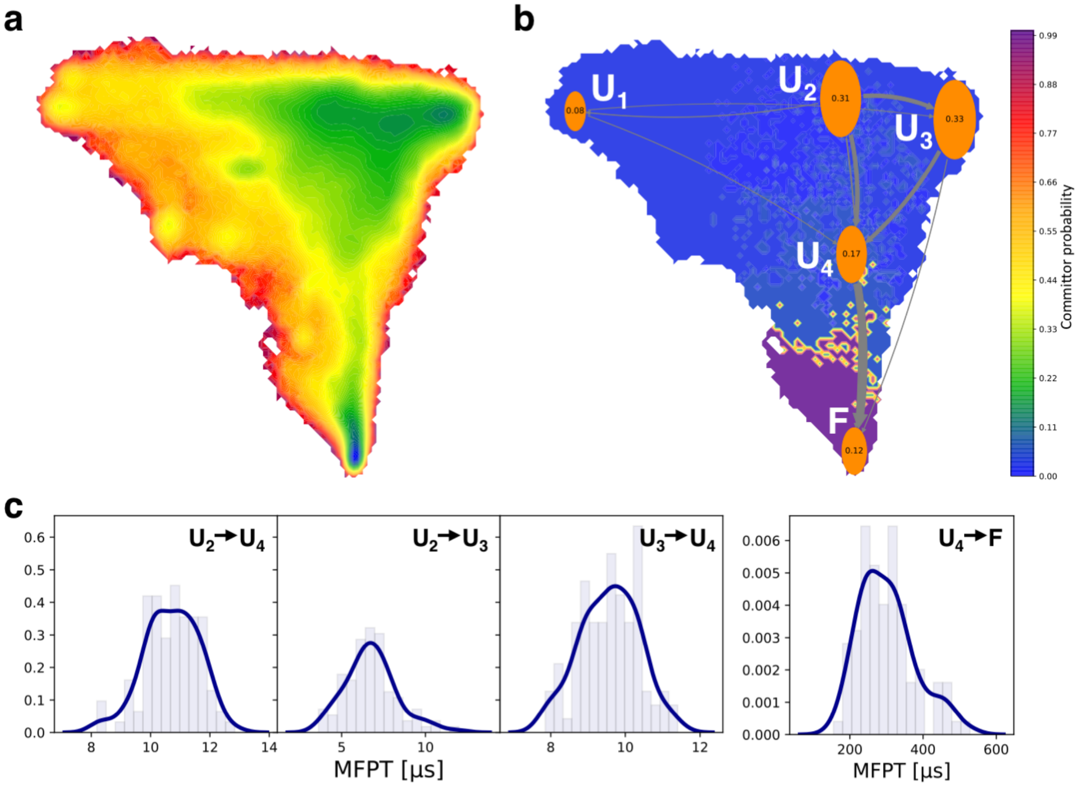


**Supplementary Figure 8. MSM analysis and MFPT estimation for protein G. a** Same free energy landscape with Fig. 4a for protein G. **b** Transition path theory analysis of the five-state hidden markov state model with stationary probability shown in orange circle. **c** Sampled mean first passage time (MFPT) for transitions of four largest flux (gray arrow) in **(b)**.

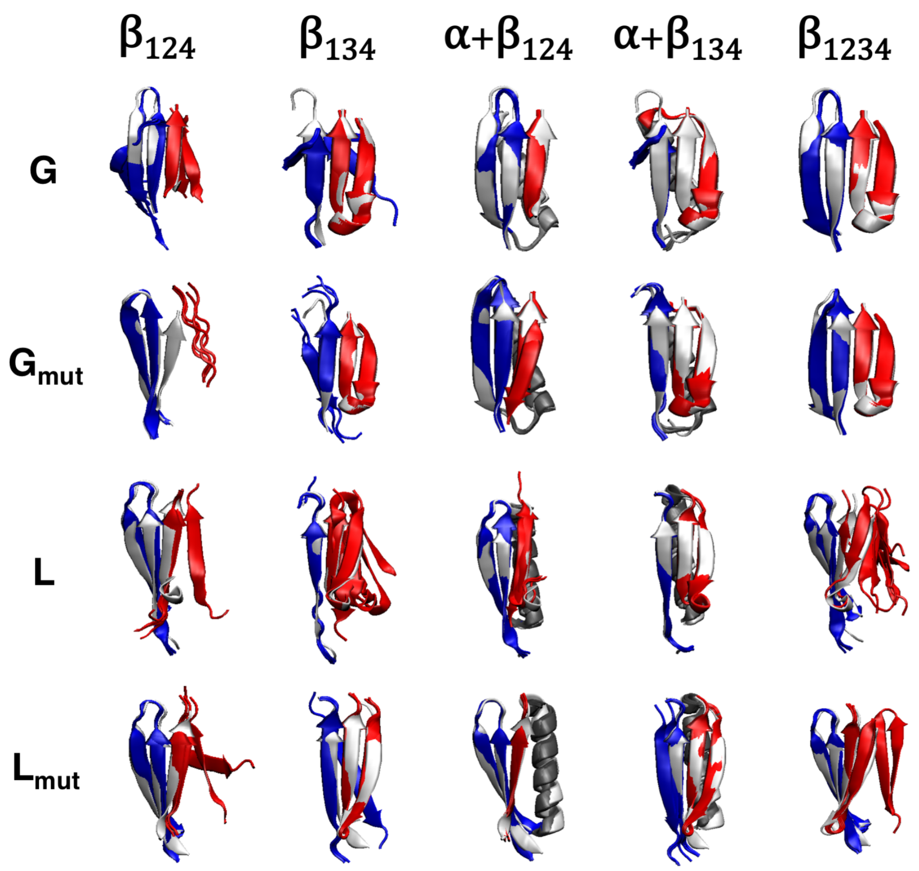


**Supplementary Figure 9. Disconnected protein fragment structure predictions from AlphaFold.** Five structure models from AF predictions with MSA for fragment $\beta_{124}$, $\beta_{134}$, $\beta_{124}$ with $\alpha$ helix, $\beta_{134}$ with $\alpha$ helix, and $\beta_{1234}$ for protein G, L and their mutants aligned with native structure (white).

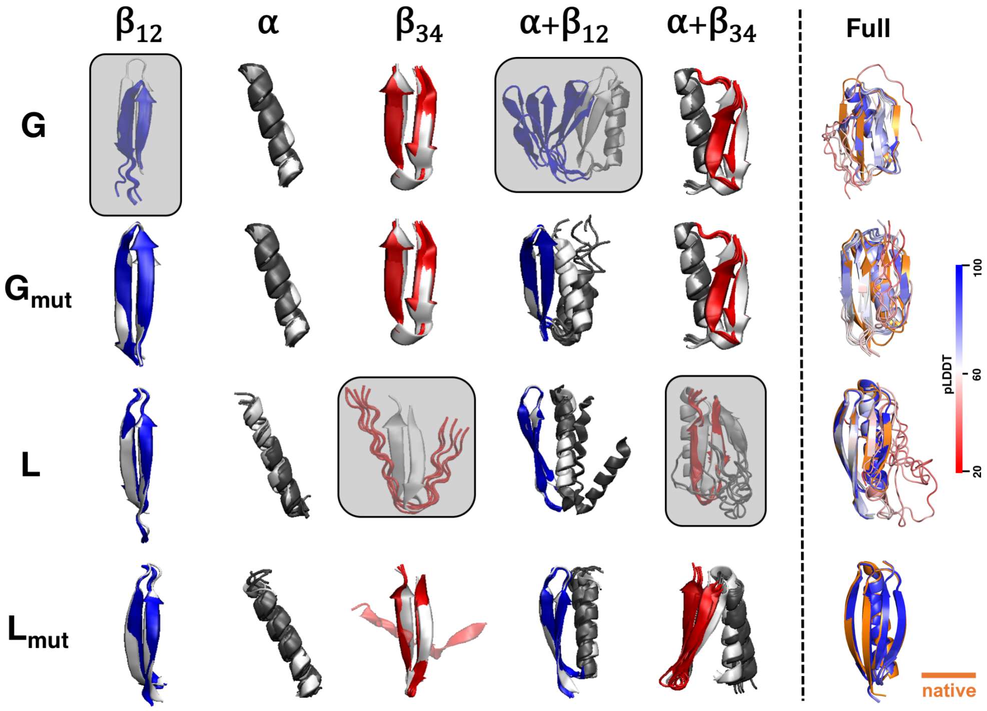


**Supplementary Figure 10. Protein fragment and full structure predictions from AlphaFold without MSA.** **Left:** Five structure models from AF predictions for fragment $\beta_{12}$ (blue), $\alpha$ helix (dark grey), $\beta_{34}$ (red), $\beta_{12}$ with $\alpha$ helix, and $\alpha$ helix with $\beta_{34}$ for protein G, L and their mutants aligned with native structure (white). **Right:** Five structure predictions colored by pLDDT score aligned with native structure (orange) without MSA for input.

### References


**1.** Roe, D. R. & Cheatham, T. E. PTRAJ and CPPTRAJ: Software for Processing and Analysis of Molecular Dynamics Trajectory Data. *J. Chem. Theory Comput.* **9**, 3084–3095 (2013).

**2.** Perez, A., Morrone, J. A., Brini, E., MacCallum, J. L. & Dill, K. A. Blind protein structure prediction using accelerated free-energy simulations. *Sci. Adv.* **2**, e1601274 (2016).

**3.** Chang, L., Perez, A. & Miranda-Quintana, R. A. Improving the analysis of biological ensembles through extended similarity measures. *Phys. Chem. Chem. Phys.* (2021).
